# A Novel Fully Automated MRI-Based Deep Learning Method for Classification of 1P/19Q Co-Deletion Status in Brain Gliomas

**DOI:** 10.1101/2020.07.15.204933

**Authors:** Chandan Ganesh Bangalore Yogananda, Bhavya R. Shah, Frank F. Yu, Marco C. Pinho, Sahil S. Nalawade, Gowtham K. Murugesan, Benjamin C. Wagner, Bruce Mickey, Toral R. Patel, Baowei Fei, Ananth J. Madhuranthakam, Joseph A. Maldjian

**Author notes:** **Corresponding Author:** Joseph A. Maldjian, MD, Department of Radiology, UT Southwestern Medical Center, 5323 Harry Hines Blvd, Dallas, Texas 75390-9178. **Authorship:** Experimental Design (C.G.B.Y., B.C.W., F.F.Y., S.S.N., G.K.M., A.J.M., J.A.M.); Implementation (C.G.B.Y., B.C.W., F.F.Y., S.S.N., G.K.M., M.C.P., A.J.M., J.A.M.); Analysis and interpretation of data (C.G.B.Y., B.R.S, S.S.N., G.K.M., F.F.Y., M.C.P.,B.M., T.R.P, B.F., A.J.M., J.A.M.); Writing of the manuscript: (C.G.B.Y., B.R.S, S.S.N., G.K.M., F.F.Y., M.C.P., B.C.W, B.M., T.R.P, B.F., A.J.M., J.A.M.).

## Abstract

**Background:** One of the most important recent discoveries in brain glioma biology has been the identification of the isocitrate dehydrogenase (IDH) mutation and 1p/19q co-deletion status as markers for therapy and prognosis. 1p/19q co-deletion is the defining genomic marker for oligodendrogliomas and confers a better prognosis and treatment response than gliomas without it. Our group has previously developed a highly accurate deep-learning network for determining IDH mutation status using T2-weighted MRI only. The purpose of this study was to develop a similar 1p/19q deep-learning classification network.

**Methods:** Multi-parametric brain MRI and corresponding genomic information were obtained for 368 subjects from The Cancer Imaging Archive (TCIA) and The Cancer Genome Atlas (TCGA). 1p/19 co-deletions were present in 130 subjects. 238 subjects were non co-deleted. A T2w image only network (1p/19q-net) was developed to perform 1p/19q co-deletion status classification and simultaneous single-label tumor segmentation using 3D-Dense-UNets. Threefold cross-validation was performed to generalize the network performance. ROC analysis was also performed. Dice-scores were computed to determine tumor segmentation accuracy.

**Results:** 1p/19q-net demonstrated a mean cross validation accuracy of 93.46% across the 3 folds (93.4%, 94.35%, and 92.62%, standard dev=0.8) in predicting 1p/19q co-deletion status with a sensitivity and specificity of 0.90 ±0.003 and 0.95 ±0.01, respectively and a mean AUC of 0.95 ±0.01. The whole tumor segmentation mean Dice-score was 0.80 ± 0.007.

**Conclusion:** We demonstrate high 1p/19q co-deletion classification accuracy using only T2-weighted MR images. This represents an important milestone toward using MRI to predict glioma histology, prognosis, and response to treatment.

**Keypoints:** 1. 1p/19 co-deletion status is an important genetic marker for gliomas. 2. We developed a non-invasive, MRI based, highly accurate deep-learning method for the determination of 1p/19q co-deletion status that only utilizes T2 weighted MR images

**IMPORTANCE OF THE STUDY:** One of the most important recent discoveries in brain glioma biology has been the identification of the isocitrate dehydrogenase (IDH) mutation and 1p/19q co-deletion status as markers for therapy and prognosis. 1p/19q co-deletion is the defining genomic marker for oligodendrogliomas and confers a better prognosis and treatment response than gliomas without it. Currently, the only reliable way to determine 1p/19q mutation status requires analysis of glioma tissue obtained either via an invasive brain biopsy or following open surgical resection. The ability to non-invasively determine 1p/19q co-deletion status has significant implications in determining therapy and predicting prognosis. We developed a highly accurate, deep learning network that utilizes only T2-weighted MR images and outperforms previously published imagebased methods. The high classification accuracy of our T2w image only network (1p/19q-net) in predicting 1p/19q co-deletion status marks an important step towards image-based stratification of brain gliomas. Imminent clinical translation is feasible because T2-weighted MR imaging is widely available and routinely performed in the assessment of gliomas.

## INTRODUCTION

Genetic profiling and molecular subtyping of glial neoplasms has revolutionized our ability to optimize therapeutic strategies and enhance prognostic accuracy. Perhaps the most compelling evidence supporting this paradigm is the 2016 revision of the World Health Organization’s (WHO) classification of gliomas which now includes genetic analysis. The impact of glioma reclassification based on molecular profiling has subsequently been studied and three genetic alterations have been extensively validated: O-6-methylguanine-DNA methyltransferase (MGMT), Isocitrate dehydrogenase (IDH), and 1p/19q co-deletion status.^1^

MGMT is a DNA repair enzyme that protects normal and glioma cells from alkylating chemotherapeutic agents. Mutations that result in methylation of the MGMT promoter result in loss of function of the enzyme and its protective effect. Mutations of IDH alter the function of the enzyme to produce D-2-hydroxyglutarate instead of α-ketoglutarate. This altered function results in increased sensitivity of the glioma to radiation and chemotherapy. Gliomas that are IDH mutated can be further divided into gliomas with or without a 1p/19q co-deletion. The 1p/19q co-deletion is defined as the combined loss of the short arm of chromosome 1 (1p) and the long arm of chromosome 19 (19q). According to the 2016 WHO classification of gliomas, an IDH mutated glioma with a 1p/19q co-deletion is classified as an oligodendroglioma, whereas an IDH mutated glioma without a 1p/19q co-deletion is classified as a diffuse astrocytoma. Oligodendrogliomas have a better prognosis when compared to diffuse astrocytomas. Additionally, even patients with an IDH-mutated *anaplastic* oligodendroglioma (WHO grade III) have a longer median overall survival than IDH-wild type, 1p/19q non co-deleted, WHO grade II astrocytomas and are more responsive to chemotherapy.^2^ Therefore, determination of 1p/19q status in IDH mutated gliomas is critical for guiding therapy and predicting prognosis. Currently, the only reliable way to determine 1p/19q mutation status requires analysis of glioma tissue obtained either via an invasive brain biopsy or following open surgical resection. These diagnostic procedures carry the burden of implicit risk. Therefore, considerable attention has been dedicated to developing non-invasive, image-based diagnostic methods.

Recent advances in deep-learning have led to a significant interest in advancing techniques for non-invasive, image-based molecular profiling of gliomas. Our group has previously demonstrated a highly-accurate, MRI-based, voxel-wise deep-learning IDH-classification network using only T2-weighted (T2w) MR images.^3^ T2w images facilitate clinical translation because they are routinely acquired, they can be obtained within 2 minutes, and high quality T2w images can even be obtained in the presence of active patient motion. Because the current standard of care for IDH mutated gliomas is heavily influenced by 1p/19q co-deletion status, the purpose of this study was to develop a highly accurate, fully automated deep-learning 3D network for 1p/19 co-deletion classification using T2-weighted images only.

## MATERIAL & METHODS

### Data and Pre-processing

Multi-parametric brain MRI data of glioma patients were obtained from the Cancer Imaging Archive (TCIA) database.^4,5^ Genomic information was provided from both the TCIA and TCGA (the cancer genome atlas) databases.^4–6^ Only pre-operative studies were used. Studies were screened for the availability of 1p/19q status and T2w image series. The final dataset of 368 subjects included 268 low grade glioma (LGG, 130 co-deleted, 138 non co-deleted) and 100 high grade glioma (HGG, all non co-deleted) subjects. TCGA subject IDs, 1p/19q co-deletion status, and tumor grade are listed in Table 1 of the supplementary data.

**Table 1.** Cross-validation results.

| **Fold**<br>**Description** | **1p/19q-net** |  |  |
| --- | --- | --- | --- |
| **Fold Number** | **% Accuracy** | **AUC** | **Dice-score** |
| Fold 1 | 93.4 | 0.9571 | 0.8151 |
| Fold 2 | 94.35 | 0.9688 | 0.8057 |
| Fold 3 | 92.62 | 0.9351 | 0.8000 |
| **AVERAGE** | **93.46 +/- 0.86** | **0.953 +/- 0.01** | **0.801 +/- 0.007** |

Tumor masks for 209 subjects were available through previous expert segmentation. ^3,7,8^ Tumor masks for the remaining 159 subjects were generated by the 3D-IDH network^3^ and validated by in-house neuro-radiologists. The tumor masks were used as ground truth for tumor segmentation in the training step. Ground truth whole tumor masks for 1p/19q co-deleted type were labelled with 1s and the ground truth tumor masks for 1p/19q non co-deleted type were labelled with 2s (Figure 1). Data preprocessing steps included (a) co-registering the T2w image to SR124 T2 template^9^ using ANTs affine registration^10^, (b) skull stripping using Brain Extraction Tool (BET)^11^ from FSL^11–14^, (c) N4BiasCorrection to remove RF inhomogeneity^15^, and (d) intensity normalization to zero-mean and unit variance. The pre-processing took less than

**Fig. 1.**
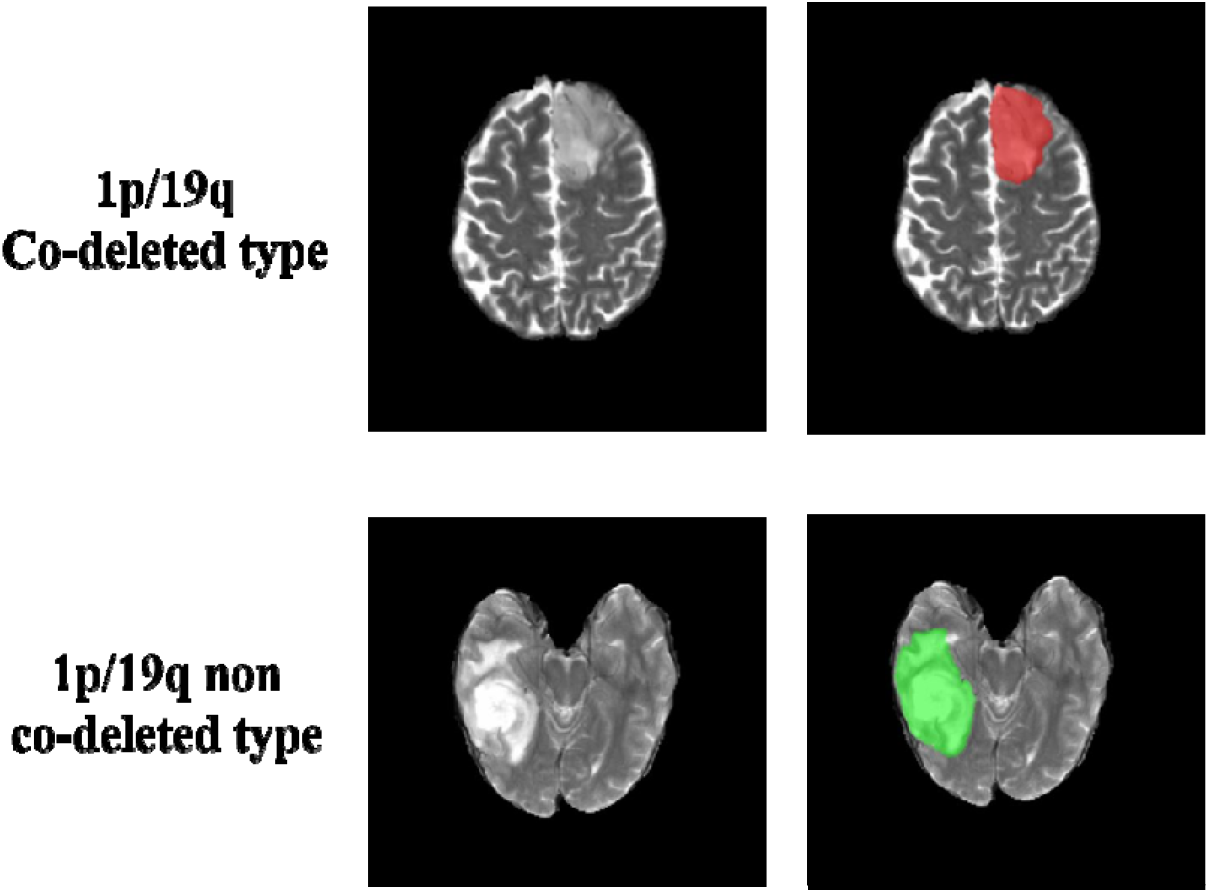
Ground truth whole tumor masks. Red voxels represent 1p/19q co-deletion status (values of 1) and green voxels represent 1p/19q non co-deletion status (values of 2). The ground truth labels have the same co-deletion status for all voxels in each tumor. 5 minutes per dataset.

### Network Details

Transfer learning was performed with the previously trained 3D-IDH network for 1p/19q classification.^3^ The decoder part of the network was fine-tuned for a voxel-wise dual-class segmentation of the whole tumor with Classes 1 and 2 representing 1p/19q co-deleted and 1p/19q non co-deleted type respectively. The schematics for the network architecture are shown in Figure 2B. A detailed description of the network is given in the supplemental material section.

**Fig. 2.**
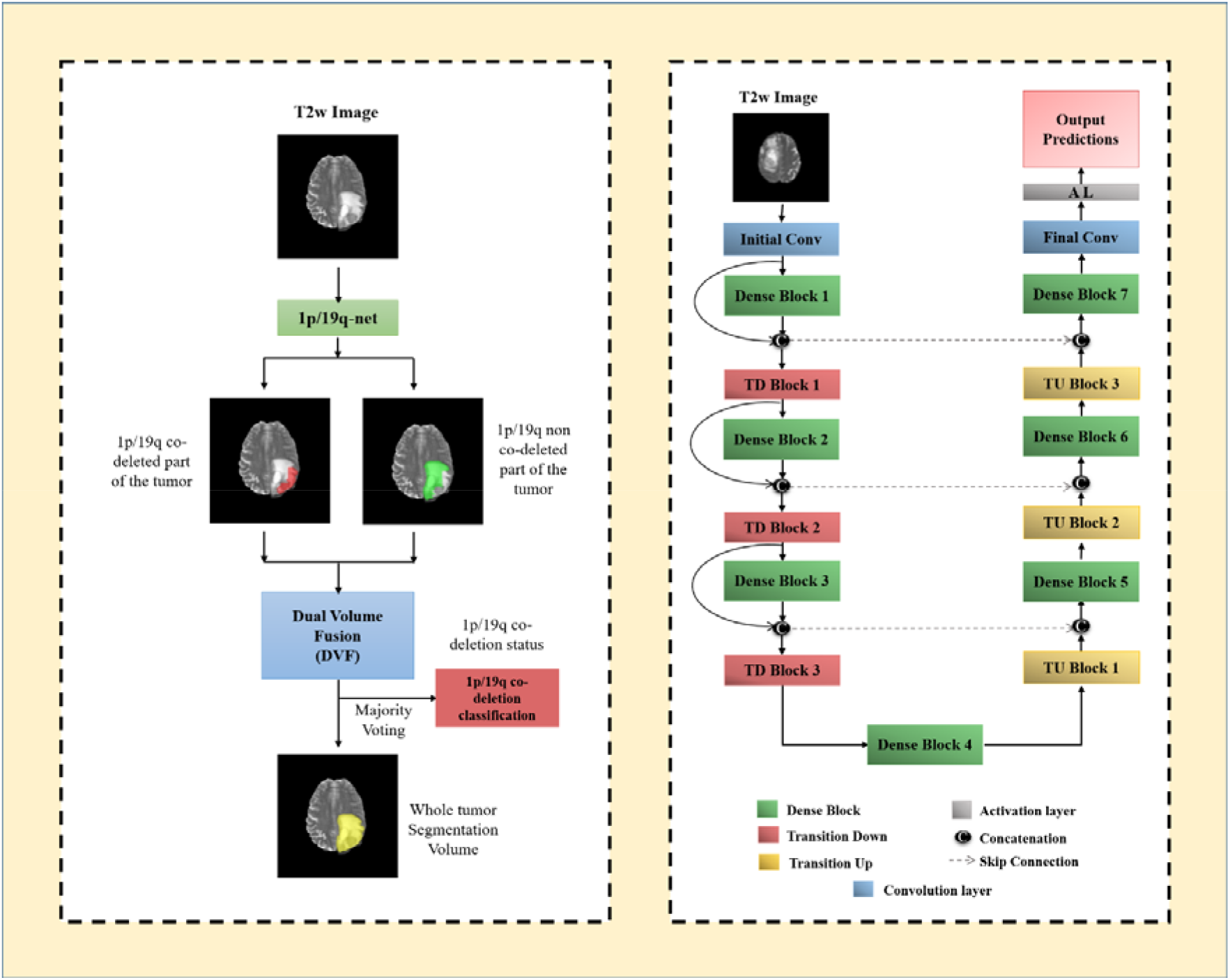
**(A) 1p/19q-net overview.** Voxel-wise classification of 1p/19q co-deletion status is performed to create 2 volumes (1p/19q co-deleted and 1p/19q non co-deleted). Volumes are combined using dual volume fusion to eliminate false positives and generate a tumor segmentation volume. Majority voting across voxels is used to determine the overall 1p/19q co-deletion status. **(B) Network architecture for 1p/19q-net.** 3D-Dense-UNets were employed with 7 dense blocks, 3 transition down blocks, and 3 transition up blocks.

### Network Implementation and Cross-validation

To generalize the reliability of the networks, a 3-fold cross-validation was performed on the 368 subjects by randomly shuffling the dataset and distributing it into 3 groups (approximately 122 subjects for each group). During each fold of the cross-validation procedure, the 3 groups were alternated between training, in-training validation and held-out testing. Group 1 had 122 subjects (43 co-deleted, 79 non co-deleted), Group 2 had 124 subjects (44 co-deleted, 80 non co-deleted), and Group 3 had 122 subjects (43 co-deleted, 79 non co-deleted). An in-training validation dataset helps the network improve its performance during training. Each fold of the cross-validation is a new training phase based on a unique combination of the 3 groups. However, network performance is only reported on the held-out testing group for each fold as it is never seen by the network. The group membership for each cross-validation fold is listed in Table 1 of the supplementary data.

Seventy-five percent overlapping patches were extracted from the training and in-training validation subjects. No patch from the same subject was mixed with the training, in-training validation or testing datasets in order to avoid the data leakage problem.^16,17^ The Data augmentation steps included vertical flipping, horizontal flipping, translation rotation, random rotation, addition of Gaussian noise, addition of salt & pepper noise and projective transformation. Additionally, all images were down-sampled by 50% and 25% (reducing the voxel resolution to 2mm x 2mm x 2mm & 4mm x 4mm x 4mm) and added to the training and validation sets. Data augmentation provided a total of approximately 300,000 patches for training and 300,000 patches for in-training validation for each fold. Networks were implemented using Keras^18^ and Tensorflow^19^ with an Adaptive Moment Estimation optimizer (Adam)^20^. The initial learning rate was set to 10^-5^ with a batch size of 15 and maximal iterations of 100. Initial parameters were chosen based on previous work with Dense-UNets using brain imaging data and semantic segmentation.^3,21,22^

1p/19q-net yields two segmentation volumes. Volume 1 provides the voxel-wise prediction of 1p/19q co-deleted tumor and Volume 2 identifies the predicted 1p/19q non codeleted tumor voxels. A single tumor segmentation map is obtained by fusing the two volumes and obtaining the largest connected component using a 3D connected component algorithm in MATLAB^(R)^. Majority voting over the voxel-wise classes of co-deleted type or non co-deleted type provided a single 1p/19q classification for each subject. Networks were implemented on Tesla V100s, P100, P40 and K80 NVIDIA-GPUs. The 1p/19q classification process developed is fully automated, and a tumor segmentation map is a natural output of the voxel-wise classification approach.

### Statistical Analysis

MATLAB^(R)^ and R were used for statistical analysis of the network’s performance. Majority voting (*i.e*. voxel-wise cutoff of 50%) was used to evaluate the accuracy of the network. The accuracy, sensitivity, specificity, positive predictive value (PPV), and negative predictive value (NPV) of the model for each fold of the cross-validation procedure were calculated using this threshold. A Receiver Operating Characteristic (ROC) curve was also generated for each fold. A Dice-score was used to evaluate the performance of the networks for tumor segmentation. The Dice-score calculates the amount of spatial overlap between the ground truth segmentation and the network segmentation.

## RESULTS

The network achieved a mean cross-validation testing accuracy of 93.46% across the 3 folds (93.4%, 94.35%, and 92.62%, standard dev=0.8). Mean cross-validation sensitivity, specificity, PPV, NPV and AUC for 1p/19q-net was 0.90 ±0.003, 0.95 ±0.01, 0.91 ±0.02, 0.95 ±0.0003 and 0.95 ±0.01 respectively. The mean cross-validation Dice-score for tumor segmentation was 0.80 ± 0.007. The network misclassified 8, 7 and 9 cases for each fold respectively (24 total out of 368 subjects). Twelve subjects were misclassified as non co-deleted, and 12 as co-deleted.

### ROC analysis

The ROC curves for each cross-validation fold for the network is provided in Figure 3. The network demonstrated very good performance curves with high sensitivities and specificities.

**Fig. 3.**
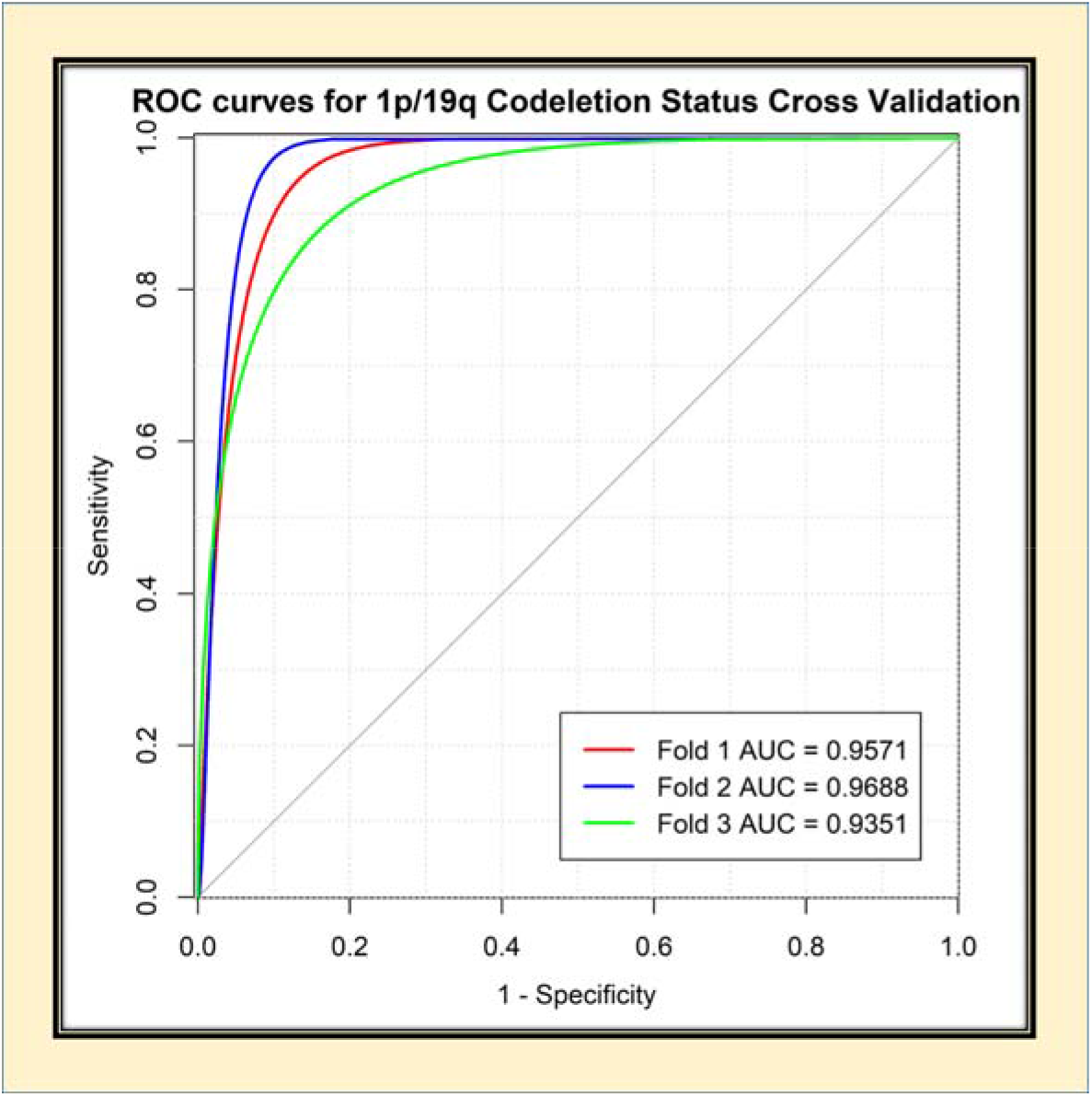
ROC analysis for 1p/19q-net. Separate curves are plottedfor each cross-validation fold along with corresponding AUC value.

### Voxel-wise classification

Since the network is a voxel-wise classifier, it performs a simultaneous tumor segmentation. Figures 4A and 4B show examples of the voxel-wise classification for a codeleted type, and non co-deleted type respectively using the network. The volume fusion procedure was effective in removing false positives to increase accuracy. This procedure improved the dice-scores by approximately 4% for the network. We also computed the voxelwise accuracy for the network. The mean voxel-wise accuracies were 85.86% ±0.01 for non codeleted type and 80.51% ±0.01 for co-deleted type.

**Fig. 4.**
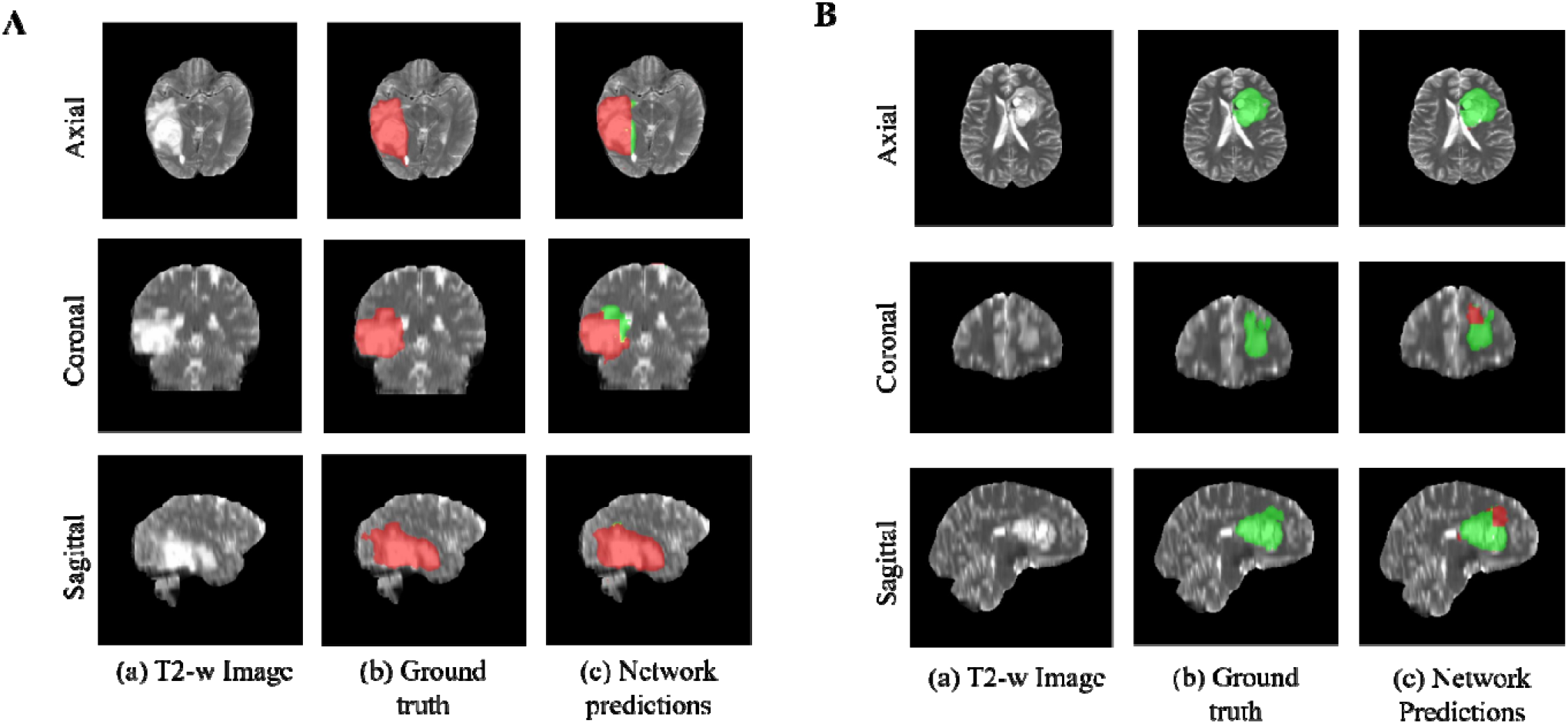
Example voxel-wise segmentation from 1p/19q-net. (A) Example for a 1p/19q co-deleted tumor. Native T2 image (a). Ground truth segmentation (b). Network output after DVF (c). Red voxels correspond to 1p/19q co-deleted class and green voxels correspond to 1p/19q non co-deleted class. (B) Example for a 1p/19q non co-deleted tumor. The sharp borders visible between co-deleted and non co-deleted type result from the patch-wise classification approach.

### Training and segmentation times

It took approximately 1 week to fine-tune the decoder portion of the network. The trained network took approximately three minutes to segment the whole tumor, and predict the 1p/19qco-deletion status for each subject.

## DISCUSSION

We developed a fully-automated, highly accurate, deep-learning network that outperforms previously reported 1p/19q co-deletion status classification algorithms.^23–26^ When comparing our T2-network with previous work, our results suggest that algorithm accuracy can be improved by using T2-weighted images only. Clinical translation becomes much simpler using only T2 weighted images because these images are routinely acquired and are robust to motion. When compared to previously published algorithms, our methodology is fully-automated. The time required for 1p/19q-net to segment the whole tumor and predict the 1p/19q co-deletion status for one subject is approximately 3 minutes on a K80, P40, P100 or V100s NVIDA-GPU.

The higher performance achieved by our network when compared to previous work is likely due to several factors. Similar to our IDH classification network we employed 3D networks whereas prior attempts at 1p/19q co-deletion status classification have relied on 2D networks.^23^ The dense connections in a 3D network architecture are advantageous because they carry information from all the previous layers to the following layers.^21^ Additionally, 3D networks are easier to train and can reduce over-fitting.^27^ As we previously reported, the Dual Volume Fusion (DVF) post-processing step helps in effectively eliminating false positives while improving the segmentation accuracy by excluding extraneous voxels not connected to the tumor. DVF improved the dice-scores by approximately 4% for the network. The 3D networks interpolate between slices to maintain inter-slice information more accurately. The network does not require extraction of pre-engineered features from the images or histopathological data^28^. Our approach also uses voxel-wise classifiers and provides a classification for each voxel in the image. This provides a simultaneous single-label tumor segmentation. Another factor that may explain the higher performance achieved by our network is that previous approaches required multi-contrast input which can be compromised due to patient motion from lengthier examination times, and the need for gadolinium contrast. High quality T2-weighted images are almost universally acquired during clinical brain tumor diagnostic evaluation. Clinically, T2w images are typically acquired within 2 minutes at the beginning of the exam and are relatively resistant to the effects of patient motion. Several of the previous 1p/19q deep learning studies were trained and tested on only low-grade gliomas achieving accuracies ranging from 65.9% - 87.7%.^23–25^ Our algorithm was trained and evaluated on a mix of high grade and low grade gliomas, which is a better representative of real-world performance and potential clinical utilization.

In the clinical setting, histologic evaluation remains the gold standard for genetic profiling of gliomas. Several different methods to detect 1p/19q co-deletion have been employed: fluorescence in-situ hybridization (FISH), array comparative genomic hybridization, multiplex ligation dependent probe amplification, and PCR-based loss of heterozygosity analysis.^29^ FISH is the most routinely performed method.^30^ FISH relies on fluorescent labeled DNA probes to directly detect chromosomal abnormalities on a tissue slide in interphase nuclei.^31^ The fraction of nuclei that demonstrate a deletion or relative deletion (in cases with polysomy) are summed and a percentage is calculated.^32^ When the percentage of “deleted” nuclei exceeds a pre-determined cut-off, the tumor is classified as 1p/19q co-deleted.^32^ A drawback of FISH is that it lacks standardized criteria for analysis of 1p/19qco-deletion status. For example, there is no consensus on what cut-off level to use when classifying co-deletion status. As a result, variability in institutional-based cut-off values can span from 20% to 70% and can affect accurate diagnosis.^32^ This limitation affects the sensitivity, specificity, PPV, and NPV of 1p/19q detection by FISH based on the cut-off value selected.^32^

There are interesting parallel considerations when studying our deep-learning method of 1p/19q determination. Our network is a voxel-wise classifier and as a result some portions within each glioma are classified as 1p/19 co-deleted while other areas are 1p/19q non co-deleted. The overall determination of 1p/19q co-deletion status is based on the majority of voxels in the tumor. Given the variability in the cut-off values for FISH detection of 1p/19q co-deletion, we performed a Youden’s statistical index analysis to determine if the optimal cut-off for our deep learning algorithm was different than majority voting (>50%). The analysis demonstrated that maximum accuracy, sensitivity, specificity, PPV, and NPV were obtained at an optimal cut-off of 50%, the same as majority voting.

The algorithm misclassified 24 cases: 12 subjects were misclassified as non co-deleted and 12 as co-deleted. Despite these misclassifications, our network achieved a mean crossvalidation testing accuracy of 93.46% which is similar to what is reported for FISH.^32^ However, our sensitivity, specificity, PPV, and NPV were significantly better than when compared to FISH.^30^ While FISH requires tissue to be obtained from an invasive procedure and subsequent tissue processing for at least 48 hours, our deep learning algorithm can segment the entire glioma and provide a 1p/19q co-deletion status in 3 minutes. The deep learning algorithm can also be fine-tuned to variations in institutional MRI scanners, while FISH analysis currently lacks standardization as mentioned above.

The limitations of our study are that deep learning studies require large amounts of data and the relative number of subjects with 1p/19q co-deletions is small. Additionally, acquisition parameters and imaging vendor platforms vary across imaging centers that contribute data.

Despite these caveats our algorithm demonstrated high 1p/19q co-deletion classification accuracy.

## CONCLUSION

We demonstrate high 1p/19q co-deletion classification accuracy using only T2-weighted MR images. This represents an important milestone toward using MRI to predict glioma histology, prognosis, and response to treatment.

## Funding

Support for this research was provided by NIH/NCI U01CA207091 (AJM, JAM).

## Disclosures

No conflicts of interest

## Acknowledgments

We thank Yin Xi, PhD, statistician for help with the ROC and AUC.

## Notes

**Funding:** This work was supported by NIH/NCI U01CA207091 (AJM, JAM).

**Conflict of Interest:** None.

### Competing Interest Statement

The authors have declared no competing interest.

