## Supplemental Table for "A Novel Fully Automated MRI-Based Deep Learning Method for Classification of 1P/19Q Co-Deletion Status in Brain Gliomas"

**SUPPLEMENTAL MATERIAL**

**FIGURES**

Supporting Figure 1: A Detailed network architecture for the 1p/19q-net. The previously trained 3D IDH network was used. The left arm of the Dense U-net (striped red box) is the encoder part of the network, the right arm of the network (blue box) is the decoder part and the dense block (yellow box) is the bottle neck block. The encoder part of the network was frozen to retain the pre-trained weights from the 3D IDH network. The bottleneck block and the decoder part of the network was fine-tuned for a dual class segmentation with class 1 representing 1p/19q co-deleted type and class 2 representing 1p/19q non co-deleted type.

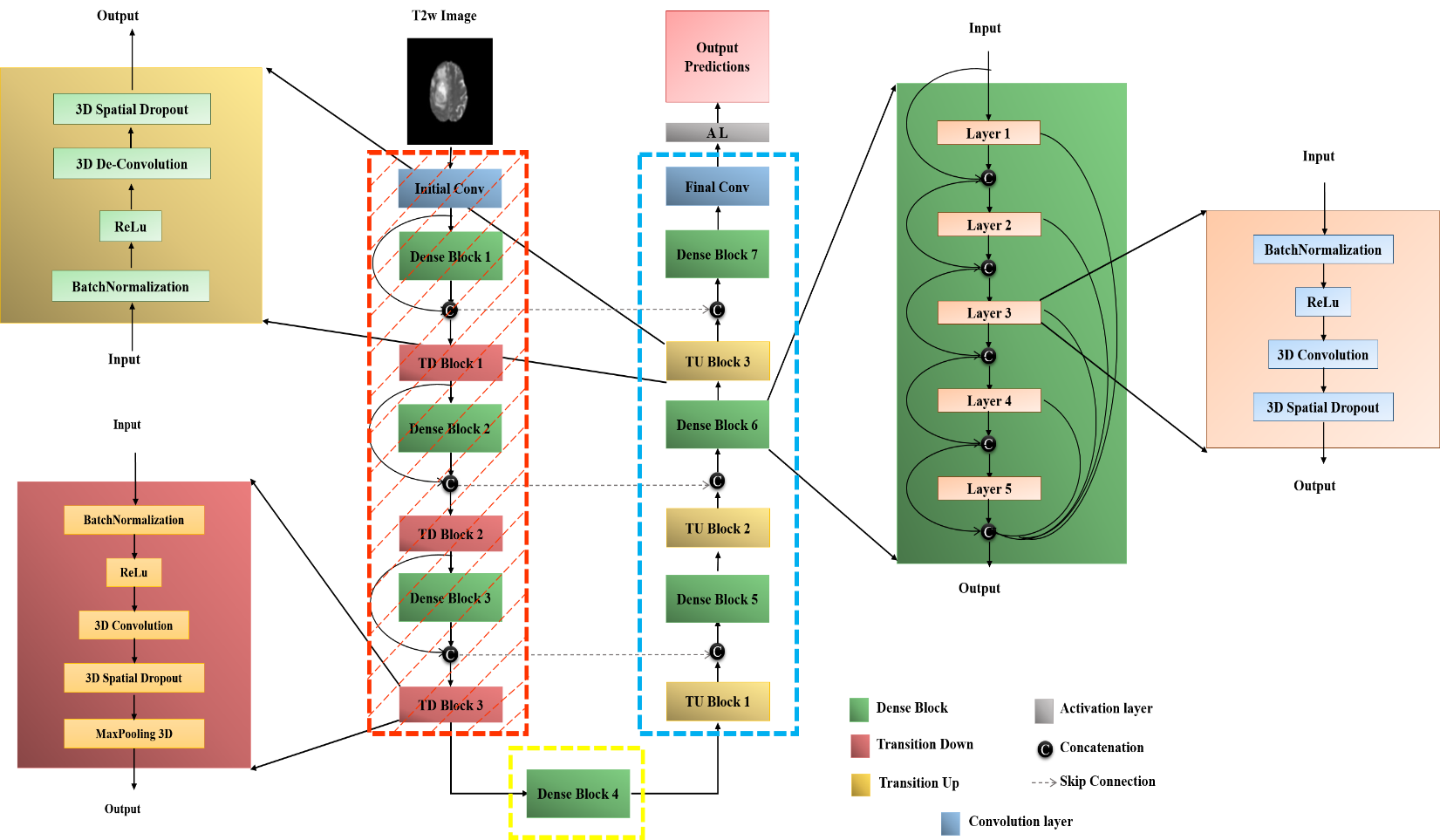

**ROC methodology**

The network output classifies voxels in the tumor as 1p/19q co-deleted or non co-deleted type. The percent of co-deleted voxels was computed for the network output for each subject in the test set by dividing the predicted co-deleted voxels by the total number of predicted voxels in each tumor. The percent co-deleted voxels can be viewed as a network output prediction likelihood of the tumor being 1p/19q co-deleted. Note that in the manuscript, majority voting (the 50% threshold) was used to determine 1p/19q co-deletion status prediction. For the ROC analysis, the percent of co-deleted voxels was sorted and used as separate thresholds (cut-points) to determine 1p/19q co-deletion status for the subjects across the test set for each new cut-point. The resulting predicted 1p/19q class membership was compared to the ground truth values to determine sensitivity (true positive rate) and 1- specificity (false positive rate) at each threshold. The resulting values were plotted using Matlab to obtain an ROC curve (true positive rate against false positive rate). Matlab routines were used to fit the curves and determine the area under the curve (AUC). This procedure was repeated for each of the 3 test folds from the cross-validation procedure for the 1p/19q-net, providing a total of 3 ROC curves from the cross-validation.

**TABLES**

Table 1: Subject wise 1p/19q co-deletion status and tumor histology

| **SUBJECT ID** | **Age** | **Gender** | **Histology** | **Grade** | **Data Collection** | **IDH**  **Status** | **IDH**  **Allele** | **1p/19q co-deletion** | **Cross-validation group** |
| --- | --- | --- | --- | --- | --- | --- | --- | --- | --- |
| TCGA-02-0003 | 50 | male | glioblastoma | G4 | HGG | WT | N/A | non-codel | 2 |
| TCGA-02-0006 | 56 | female | glioblastoma | G4 | HGG | WT | N/A | non-codel | 3 |
| TCGA-02-0009 | 61 | female | glioblastoma | G4 | HGG | WT | N/A | non-codel | 1 |
| TCGA-02-0011 | 18 | female | glioblastoma | G4 | HGG | WT | N/A | non-codel | 1 |
| TCGA-02-0027 | 33 | female | glioblastoma | G4 | HGG | WT | N/A | non-codel | 1 |
| TCGA-02-0033 | 54 | male | glioblastoma | G4 | HGG | WT | N/A | non-codel | 1 |
| TCGA-02-0034 | 60 | male | glioblastoma | G4 | HGG | WT | N/A | non-codel | 3 |
| TCGA-02-0037 | 74 | female | glioblastoma | G4 | HGG | WT | N/A | non-codel | 2 |
| TCGA-02-0046 | 61 | male | glioblastoma | G4 | HGG | WT | N/A | non-codel | 2 |
| TCGA-02-0047 | 78 | male | glioblastoma | G4 | HGG | WT | N/A | non-codel | 3 |
| TCGA-02-0048 | 80 | male | glioblastoma | G4 | HGG | WT | N/A | non-codel | 3 |
| TCGA-02-0054 | 44 | female | glioblastoma | G4 | HGG | WT | N/A | non-codel | 1 |
| TCGA-02-0060 | 66 | female | glioblastoma | G4 | HGG | WT | N/A | non-codel | 3 |
| TCGA-02-0064 | 50 | male | glioblastoma | G4 | HGG | WT | N/A | non-codel | 3 |
| TCGA-02-0068 | 57 | male | glioblastoma | G4 | HGG | WT | N/A | non-codel | 2 |
| TCGA-02-0069 | 31 | female | glioblastoma | G4 | HGG | WT | N/A | non-codel | 1 |
| TCGA-02-0070 | 70 | male | glioblastoma | G4 | HGG | WT | N/A | non-codel | 2 |
| TCGA-02-0075 | 63 | male | glioblastoma | G4 | HGG | WT | N/A | non-codel | 2 |
| TCGA-02-0085 | 63 | female | glioblastoma | G4 | HGG | WT | N/A | non-codel | 3 |
| TCGA-02-0086 | 45 | female | glioblastoma | G4 | HGG | WT | N/A | non-codel | 1 |
| TCGA-02-0102 | 42 | male | glioblastoma | G4 | HGG | WT | N/A | non-codel | 2 |
| TCGA-06-0119 | 81 | female | glioblastoma | G4 | HGG | WT | N/A | non-codel | 2 |
| TCGA-06-0122 | 84 | female | glioblastoma | G4 | HGG | WT | N/A | non-codel | 3 |
| TCGA-06-0127 | 67 | male | glioblastoma | G4 | HGG | WT | N/A | non-codel | 1 |
| TCGA-06-0128 | 66 | male | glioblastoma | G4 | HGG | Mutant | IDH1 | non-codel | 1 |
| TCGA-06-0129 | 30 | male | glioblastoma | G4 | HGG | Mutant | IDH1 | non-codel | 3 |
| TCGA-06-0132 | 49 | male | glioblastoma | G4 | HGG | WT | N/A | non-codel | 2 |
| TCGA-06-0133 | 64 | male | glioblastoma | G4 | HGG | WT | N/A | non-codel | 3 |
| TCGA-06-0137 | 63 | female | glioblastoma | G4 | HGG | WT | N/A | non-codel | 2 |
| TCGA-06-0138 | 43 | male | glioblastoma | G4 | HGG | WT | N/A | non-codel | 2 |
| TCGA-06-0139 | 40 | male | glioblastoma | G4 | HGG | WT | N/A | non-codel | 2 |
| TCGA-06-0142 | 81 | male | glioblastoma | G4 | HGG | WT | N/A | non-codel | 3 |
| TCGA-06-0143 | 58 | male | glioblastoma | G4 | HGG | WT | N/A | non-codel | 3 |
| TCGA-06-0145 | 53 | female | glioblastoma | G4 | HGG | WT | N/A | non-codel | 3 |
| TCGA-06-0147 | 51 | female | glioblastoma | G4 | HGG | WT | N/A | non-codel | 1 |
| TCGA-06-0154 | 54 | male | glioblastoma | G4 | HGG | WT | N/A | non-codel | 2 |
| TCGA-06-0157 | 63 | female | glioblastoma | G4 | HGG | WT | N/A | non-codel | 1 |
| TCGA-06-0158 | 73 | male | glioblastoma | G4 | HGG | WT | N/A | non-codel | 3 |
| TCGA-06-0166 | 51 | male | glioblastoma | G4 | HGG | WT | N/A | non-codel | 1 |
| TCGA-06-0168 | 59 | female | glioblastoma | G4 | HGG | WT | N/A | non-codel | 1 |
| TCGA-06-0174 | 54 | male | glioblastoma | G4 | HGG | WT | N/A | non-codel | 2 |
| TCGA-06-0176 | 34 | male | glioblastoma | G4 | HGG | WT | N/A | non-codel | 2 |
| TCGA-06-0184 | 63 | male | glioblastoma | G4 | HGG | WT | N/A | non-codel | 3 |
| TCGA-06-0185 | 54 | male | glioblastoma | G4 | HGG | WT | N/A | non-codel | 1 |
| TCGA-06-0187 | 69 | male | glioblastoma | G4 | HGG | WT | N/A | non-codel | 3 |
| TCGA-06-0188 | 71 | male | glioblastoma | G4 | HGG | WT | N/A | non-codel | 3 |
| TCGA-06-0189 | 55 | male | glioblastoma | G4 | HGG | WT | N/A | non-codel | 1 |
| TCGA-06-0190 | 62 | male | glioblastoma | G4 | HGG | WT | N/A | non-codel | 2 |
| TCGA-06-0192 | 58 | male | glioblastoma | G4 | HGG | WT | N/A | non-codel | 1 |
| TCGA-06-0213 | 55 | female | glioblastoma | G4 | HGG | WT | N/A | non-codel | 2 |
| TCGA-06-0237 | 75 | female | glioblastoma | G4 | HGG | WT | N/A | non-codel | 2 |
| TCGA-06-0238 | 46 | male | glioblastoma | G4 | HGG | WT | N/A | non-codel | 2 |
| TCGA-06-0241 | 65 | female | glioblastoma | G4 | HGG | WT | N/A | non-codel | 1 |
| TCGA-06-0644 | 71 | male | glioblastoma | G4 | HGG | WT | N/A | non-codel | 3 |
| TCGA-06-0645 | 55 | female | glioblastoma | G4 | HGG | WT | N/A | non-codel | 1 |
| TCGA-06-0646 | 60 | male | glioblastoma | G4 | HGG | WT | N/A | non-codel | 1 |
| TCGA-06-0648 | 77 | male | glioblastoma | G4 | HGG | WT | N/A | non-codel | 1 |
| TCGA-06-0649 | 73 | female | glioblastoma | G4 | HGG | WT | N/A | non-codel | 2 |
| TCGA-06-1806 | 47 | male | glioblastoma | G4 | HGG | WT | N/A | non-codel | 2 |
| TCGA-06-2570 | 21 | female | glioblastoma | G4 | HGG | Mutant | IDH1 | non-codel | 1 |
| TCGA-06-5408 | 54 | female | glioblastoma | G4 | HGG | WT | N/A | non-codel | 2 |
| TCGA-06-5412 | 78 | female | glioblastoma | G4 | HGG | WT | N/A | non-codel | 3 |
| TCGA-06-5413 | 67 | male | glioblastoma | G4 | HGG | WT | N/A | non-codel | 1 |
| TCGA-06-6389 | 49 | female | glioblastoma | G4 | HGG | Mutant | IDH1 | non-codel | 2 |
| TCGA-08-0390 | 69 | male | glioblastoma | G4 | HGG | WT | N/A | non-codel | 3 |
| TCGA-12-0616 | 36 | female | glioblastoma | G4 | HGG | WT | N/A | non-codel | 1 |
| TCGA-12-0829 | 75 | male | glioblastoma | G4 | HGG | WT | N/A | non-codel | 3 |
| TCGA-12-1093 | 66 | female | glioblastoma | G4 | HGG | WT | N/A | non-codel | 2 |
| TCGA-12-1598 | 75 | female | glioblastoma | G4 | HGG | WT | N/A | non-codel | 2 |
| TCGA-12-1602 | 58 | male | glioblastoma | G4 | HGG | WT | N/A | non-codel | 2 |
| TCGA-12-3650 | 46 | male | glioblastoma | G4 | HGG | WT | N/A | non-codel | 3 |
| TCGA-14-0789 | 54 | male | glioblastoma | G4 | HGG | WT | N/A | non-codel | 3 |
| TCGA-14-1456 | 23 | male | glioblastoma | G4 | HGG | Mutant | IDH1 | non-codel | 2 |
| TCGA-14-1794 | 59 | male | glioblastoma | G4 | HGG | WT | N/A | non-codel | 3 |
| TCGA-14-1829 | 57 | male | glioblastoma | G4 | HGG | WT | N/A | non-codel | 1 |
| TCGA-14-3477 | 38 | female | glioblastoma | G4 | HGG | WT | N/A | non-codel | 2 |
| TCGA-19-1388 | 58 | male | glioblastoma | G4 | HGG | WT | N/A | non-codel | 1 |
| TCGA-19-1390 | 63 | female | glioblastoma | G4 | HGG | WT | N/A | non-codel | 1 |
| TCGA-19-1789 | 69 | female | glioblastoma | G4 | HGG | WT | N/A | non-codel | 2 |
| TCGA-19-2624 | 51 | male | glioblastoma | G4 | HGG | WT | N/A | non-codel | 3 |
| TCGA-19-2631 | 74 | female | glioblastoma | G4 | HGG | WT | N/A | non-codel | 1 |
| TCGA-19-5954 | 72 | female | glioblastoma | G4 | HGG | WT | N/A | non-codel | 3 |
| TCGA-19-5958 | 56 | male | glioblastoma | G4 | HGG | WT | N/A | non-codel | 1 |
| TCGA-27-1835 | 53 | female | glioblastoma | G4 | HGG | WT | N/A | non-codel | 3 |
| TCGA-27-1838 | 59 | female | glioblastoma | G4 | HGG | WT | N/A | non-codel | 2 |
| TCGA-76-4926 | 68 | male | glioblastoma | G4 | HGG | WT | N/A | non-codel | 3 |
| TCGA-76-4934 | 66 | female | glioblastoma | G4 | HGG | WT | N/A | non-codel | 2 |
| TCGA-76-4935 | 52 | female | glioblastoma | G4 | HGG | WT | N/A | non-codel | 3 |
| TCGA-76-6191 | 57 | male | glioblastoma | G4 | HGG | WT | N/A | non-codel | 3 |
| TCGA-76-6192 | 74 | male | glioblastoma | G4 | HGG | WT | N/A | non-codel | 1 |
| TCGA-76-6193 | 78 | male | glioblastoma | G4 | HGG | WT | N/A | non-codel | 3 |
| TCGA-76-6280 | 57 | male | glioblastoma | G4 | HGG | WT | N/A | non-codel | 2 |
| TCGA-76-6282 | 63 | male | glioblastoma | G4 | HGG | WT | N/A | non-codel | 2 |
| TCGA-76-6285 | 64 | female | glioblastoma | G4 | HGG | WT | N/A | non-codel | 2 |
| TCGA-76-6656 | 66 | male | glioblastoma | G4 | HGG | WT | N/A | non-codel | 3 |
| TCGA-76-6657 | 74 | male | glioblastoma | G4 | HGG | WT | N/A | non-codel | 1 |
| TCGA-76-6661 | 54 | male | glioblastoma | G4 | HGG | WT | N/A | non-codel | 1 |
| TCGA-76-6662 | 58 | male | glioblastoma | G4 | HGG | WT | N/A | non-codel | 1 |
| TCGA-76-6663 | 44 | female | glioblastoma | G4 | HGG | WT | N/A | non-codel | 1 |
| TCGA-76-6664 | 49 | female | glioblastoma | G4 | HGG | WT | N/A | non-codel | 2 |
| TCGA-CS-4941 | 67 | male | astrocytoma | G3 | LGG | WT | N/A | non-codel | 2 |
| TCGA-CS-4942 | 44 | female | astrocytoma | G3 | LGG | Mutant | IDH1 | non-codel | 2 |
| TCGA-CS-4943 | 37 | male | astrocytoma | G3 | LGG | Mutant | IDH1 | non-codel | 2 |
| TCGA-CS-4944 | 50 | male | astrocytoma | G2 | LGG | Mutant | IDH1 | non-codel | 3 |
| TCGA-CS-5393 | 39 | male | astrocytoma | G3 | LGG | Mutant | IDH1 | non-codel | 3 |
| TCGA-CS-5395 | 43 | male | oligodendroglioma | G2 | LGG | WT | N/A | non-codel | 2 |
| TCGA-CS-5396 | 53 | female | oligodendroglioma | G3 | LGG | Mutant | IDH1 | codel | 2 |
| TCGA-CS-5397 | 54 | female | astrocytoma | G3 | LGG | WT | N/A | non-codel | 3 |
| TCGA-CS-6186 | 58 | male | oligoastrocytoma | G3 | LGG | WT | N/A | non-codel | 1 |
| TCGA-CS-6188 | 48 | male | astrocytoma | G3 | LGG | WT | N/A | non-codel | 1 |
| TCGA-CS-6290 | 31 | male | astrocytoma | G3 | LGG | Mutant | IDH1 | non-codel | 3 |
| TCGA-CS-6665 | 51 | female | astrocytoma | G3 | LGG | Mutant | IDH1 | non-codel | 2 |
| TCGA-CS-6666 | 22 | male | astrocytoma | G3 | LGG | Mutant | IDH1 | non-codel | 3 |
| TCGA-CS-6667 | 39 | female | astrocytoma | G2 | LGG | Mutant | IDH1 | non-codel | 1 |
| TCGA-CS-6668 | 57 | female | oligodendroglioma | G2 | LGG | Mutant | IDH1 | codel | 2 |
| TCGA-CS-6669 | 26 | female | oligodendroglioma | G2 | LGG | WT | N/A | non-codel | 1 |
| TCGA-DU-5849 | 48 | male | oligodendroglioma | G2 | LGG | Mutant | IDH1 | codel | 1 |
| TCGA-DU-5851 | 40 | female | oligoastrocytoma | G3 | LGG | Mutant | IDH1 | non-codel | 3 |
| TCGA-DU-5852 | 61 | female | oligoastrocytoma | G3 | LGG | WT | N/A | non-codel | 1 |
| TCGA-DU-5853 | 29 | male | oligoastrocytoma | G2 | LGG | Mutant | IDH1 | non-codel | 3 |
| TCGA-DU-5854 | 57 | female | astrocytoma | G3 | LGG | WT | N/A | non-codel | 2 |
| TCGA-DU-5855 | 49 | female | oligoastrocytoma | G3 | LGG | Mutant | IDH1 | non-codel | 1 |
| TCGA-DU-5871 | 37 | female | oligoastrocytoma | G2 | LGG | Mutant | IDH1 | non-codel | 2 |
| TCGA-DU-5872 | 43 | female | oligoastrocytoma | G2 | LGG | Mutant | IDH1 | non-codel | 1 |
| TCGA-DU-5874 | 62 | female | oligodendroglioma | G2 | LGG | Mutant | IDH1 | codel | 2 |
| TCGA-DU-6395 | 31 | male | oligoastrocytoma | G2 | LGG | Mutant | IDH1 | non-codel | 1 |
| TCGA-DU-6397 | 45 | male | oligodendroglioma | G3 | LGG | Mutant | IDH1 | codel | 2 |
| TCGA-DU-6399 | 54 | male | oligodendroglioma | G2 | LGG | Mutant | IDH1 | non-codel | 2 |
| TCGA-DU-6400 | 66 | female | oligodendroglioma | G2 | LGG | Mutant | IDH1 | codel | 1 |
| TCGA-DU-6401 | 31 | female | oligodendroglioma | G2 | LGG | Mutant | IDH1 | non-codel | 3 |
| TCGA-DU-6404 | 24 | female | oligodendroglioma | G3 | LGG | WT | N/A | non-codel | 3 |
| TCGA-DU-6405 | 51 | female | astrocytoma | G3 | LGG | WT | N/A | non-codel | 3 |
| TCGA-DU-6407 | 35 | female | oligodendroglioma | G2 | LGG | Mutant | IDH1 | non-codel | 3 |
| TCGA-DU-6408 | 23 | female | oligodendroglioma | G3 | LGG | Mutant | IDH1 | non-codel | 3 |
| TCGA-DU-7008 | 41 | female | oligodendroglioma | G2 | LGG | Mutant | IDH1 | non-codel | 2 |
| TCGA-DU-7010 | 58 | female | astrocytoma | G3 | LGG | Mutant | IDH1 | non-codel | 1 |
| TCGA-DU-7015 | 41 | female | oligodendroglioma | G2 | LGG | Mutant | IDH1 | non-codel | 3 |
| TCGA-DU-7018 | 57 | female | oligodendroglioma | G3 | LGG | Mutant | IDH1 | codel | 1 |
| TCGA-DU-7019 | 39 | male | oligoastrocytoma | G3 | LGG | Mutant | IDH1 | non-codel | 2 |
| TCGA-DU-7294 | 53 | female | oligodendroglioma | G2 | LGG | Mutant | IDH1 | codel | 2 |
| TCGA-DU-7298 | 38 | female | astrocytoma | G3 | LGG | Mutant | IDH1 | non-codel | 1 |
| TCGA-DU-7299 | 33 | male | astrocytoma | G3 | LGG | Mutant | IDH1 | non-codel | 1 |
| TCGA-DU-7300 | 53 | female | oligodendroglioma | G3 | LGG | Mutant | IDH1 | codel | 3 |
| TCGA-DU-7301 | 53 | male | oligodendroglioma | G2 | LGG | Mutant | IDH1 | non-codel | 2 |
| TCGA-DU-7302 | 48 | female | oligodendroglioma | G3 | LGG | Mutant | IDH1 | codel | 3 |
| TCGA-DU-7304 | 43 | male | oligoastrocytoma | G3 | LGG | Mutant | IDH1 | non-codel | 1 |
| TCGA-DU-7306 | 67 | male | oligoastrocytoma | G2 | LGG | Mutant | IDH1 | non-codel | 1 |
| TCGA-DU-7309 | 41 | female | oligodendroglioma | G3 | LGG | Mutant | IDH2 | non-codel | 3 |
| TCGA-DU-8162 | 61 | female | oligoastrocytoma | G3 | LGG | WT | N/A | non-codel | 2 |
| TCGA-DU-8163 | 29 | male | oligoastrocytoma | G3 | LGG | Mutant | IDH1 | non-codel | 3 |
| TCGA-DU-8164 | 51 | male | oligodendroglioma | G2 | LGG | Mutant | IDH1 | codel | 1 |
| TCGA-DU-8165 | 60 | female | oligodendroglioma | G3 | LGG | WT | N/A | non-codel | 3 |
| TCGA-DU-8166 | 29 | female | oligoastrocytoma | G2 | LGG | Mutant | IDH1 | non-codel | 2 |
| TCGA-DU-8167 | 69 | female | oligoastrocytoma | G2 | LGG | Mutant | IDH1 | non-codel | 2 |
| TCGA-DU-8168 | 55 | female | oligodendroglioma | G3 | LGG | Mutant | IDH1 | codel | 3 |
| TCGA-DU-A5TP | 33 | male | astrocytoma | G3 | LGG | Mutant | IDH1 | non-codel | 2 |
| TCGA-DU-A5TR | 51 | male | oligoastrocytoma | G2 | LGG | Mutant | IDH1 | non-codel | 2 |
| TCGA-DU-A5TS | 42 | male | oligodendroglioma | G2 | LGG | Mutant | IDH1 | non-codel | 1 |
| TCGA-DU-A5TT | 70 | male | oligodendroglioma | G3 | LGG | WT | N/A | non-codel | 2 |
| TCGA-DU-A5TU | 62 | female | astrocytoma | G2 | LGG | Mutant | IDH1 | non-codel | 2 |
| TCGA-DU-A5TW | 33 | female | astrocytoma | G3 | LGG | Mutant | IDH1 | non-codel | 2 |
| TCGA-DU-A5TY | 46 | female | astrocytoma | G3 | LGG | WT | N/A | non-codel | 2 |
| TCGA-DU-A6S2 | 37 | female | oligodendroglioma | G2 | LGG | Mutant | IDH1 | codel | 2 |
| TCGA-DU-A6S3 | 60 | male | oligodendroglioma | G2 | LGG | Mutant | IDH1 | codel | 3 |
| TCGA-DU-A6S6 | 35 | female | oligoastrocytoma | G2 | LGG | Mutant | IDH1 | codel | 1 |
| TCGA-DU-A6S7 | 27 | female | astrocytoma | G3 | LGG | Mutant | IDH1 | non-codel | 3 |
| TCGA-DU-A6S8 | 74 | female | oligodendroglioma | G3 | LGG | Mutant | IDH1 | codel | 1 |
| TCGA-FG-5964 | 62 | male | oligodendroglioma | G2 | LGG | Mutant | IDH1 | codel | 2 |
| TCGA-FG-6688 | 59 | female | astrocytoma | G3 | LGG | WT | N/A | non-codel | 1 |
| TCGA-FG-6689 | 30 | male | astrocytoma | G2 | LGG | Mutant | IDH1 | non-codel | 1 |
| TCGA-FG-6690 | 70 | male | oligodendroglioma | G2 | LGG | Mutant | IDH1 | non-codel | 3 |
| TCGA-FG-6691 | 23 | female | astrocytoma | G2 | LGG | Mutant | IDH1 | non-codel | 1 |
| TCGA-FG-6692 | 63 | male | oligodendroglioma | G3 | LGG | WT | N/A | non-codel | 3 |
| TCGA-FG-7634 | 28 | male | oligodendroglioma | G2 | LGG | Mutant | IDH1 | codel | 2 |
| TCGA-FG-7643 | 49 | female | oligoastrocytoma | G2 | LGG | WT | N/A | non-codel | 1 |
| TCGA-FG-8189 | 33 | female | oligodendroglioma | G2 | LGG | Mutant | IDH2 | non-codel | 2 |
| TCGA-FG-A4MT | 27 | female | oligodendroglioma | G2 | LGG | Mutant | IDH1 | non-codel | 2 |
| TCGA-FG-A6IZ | 60 | male | oligodendroglioma | G2 | LGG | Mutant | IDH1 | codel | 3 |
| TCGA-FG-A713 | 74 | female | oligoastrocytoma | G2 | LGG | Mutant | IDH1 | codel | 1 |
| TCGA-HT-7473 | 28 | male | oligoastrocytoma | G2 | LGG | Mutant | IDH1 | non-codel | 3 |
| TCGA-HT-7475 | 67 | male | oligoastrocytoma | G3 | LGG | Mutant | IDH1 | non-codel | 3 |
| TCGA-HT-7602 | 21 | male | oligodendroglioma | G2 | LGG | Mutant | IDH1 | non-codel | 3 |
| TCGA-HT-7604 | 50 | male | astrocytoma | G2 | LGG | Mutant | IDH1 | non-codel | 1 |
| TCGA-HT-7605 | 38 | male | oligodendroglioma | G2 | LGG | Mutant | IDH1 | codel | 3 |
| TCGA-HT-7608 | 61 | male | oligoastrocytoma | G2 | LGG | Mutant | IDH1 | codel | 3 |
| TCGA-HT-7616 | 75 | male | oligodendroglioma | G3 | LGG | Mutant | IDH1 | codel | 2 |
| TCGA-HT-7680 | 32 | female | astrocytoma | G2 | LGG | WT | N/A | non-codel | 1 |
| TCGA-HT-7686 | 29 | female | astrocytoma | G3 | LGG | Mutant | IDH1 | non-codel | 2 |
| TCGA-HT-7690 | 29 | male | oligoastrocytoma | G3 | LGG | Mutant | IDH1 | non-codel | 3 |
| TCGA-HT-7692 | 43 | male | oligoastrocytoma | G2 | LGG | Mutant | IDH1 | codel | 3 |
| TCGA-HT-7693 | 51 | female | oligodendroglioma | G2 | LGG | Mutant | IDH1 | non-codel | 1 |
| TCGA-HT-7694 | 60 | male | oligodendroglioma | G3 | LGG | Mutant | IDH1 | codel | 1 |
| TCGA-HT-7855 | 39 | male | astrocytoma | G3 | LGG | Mutant | IDH1 | non-codel | 1 |
| TCGA-HT-7856 | 35 | male | oligodendroglioma | G3 | LGG | Mutant | IDH2 | codel | 1 |
| TCGA-HT-7860 | 60 | female | astrocytoma | G3 | LGG | WT | N/A | non-codel | 1 |
| TCGA-HT-7874 | 41 | female | oligodendroglioma | G3 | LGG | Mutant | IDH1 | codel | 1 |
| TCGA-HT-7879 | 31 | male | oligoastrocytoma | G3 | LGG | Mutant | IDH1 | non-codel | 3 |
| TCGA-HT-7882 | 66 | male | oligodendroglioma | G3 | LGG | WT | N/A | non-codel | 2 |
| TCGA-HT-7884 | 44 | female | astrocytoma | G2 | LGG | Mutant | IDH1 | non-codel | 1 |
| TCGA-HT-8018 | 40 | female | oligoastrocytoma | G2 | LGG | Mutant | IDH1 | non-codel | 1 |
| TCGA-HT-8105 | 54 | male | oligodendroglioma | G3 | LGG | Mutant | IDH1 | codel | 3 |
| TCGA-HT-8106 | 53 | male | astrocytoma | G3 | LGG | Mutant | IDH1 | non-codel | 3 |
| TCGA-HT-8107 | 62 | male | oligodendroglioma | G2 | LGG | WT | N/A | non-codel | 3 |
| TCGA-HT-8111 | 32 | male | oligoastrocytoma | G3 | LGG | Mutant | IDH1 | non-codel | 2 |
| TCGA-HT-8113 | 49 | female | oligodendroglioma | G2 | LGG | Mutant | IDH2 | non-codel | 2 |
| TCGA-HT-8114 | 36 | male | oligoastrocytoma | G3 | LGG | Mutant | IDH1 | non-codel | 2 |
| TCGA-HT-8563 | 30 | female | astrocytoma | G3 | LGG | Mutant | IDH1 | non-codel | 3 |
| TCGA-HT-A5RC | 70 | female | astrocytoma | G3 | LGG | WT | N/A | non-codel | 3 |
| TCGA-HT-A61A | 20 | female | oligodendroglioma | G2 | LGG | Mutant | IDH1 | non-codel | 1 |
| LGG-104 | NaN | NaN | Oligoastrocytoma | G3 | LGG | NaN | NaN | codel | 1 |
| LGG-203 | NaN | NaN | Astrocytoma | G3 | LGG | NaN | NaN | non-codel | 1 |
| LGG-210 | NaN | NaN | Oligoastrocytoma | G2 | LGG | NaN | NaN | non-codel | 1 |
| LGG-216 | NaN | NaN | Oligoastrocytoma | G2 | LGG | NaN | NaN | codel | 2 |
| LGG-218 | NaN | NaN | Oligodendroglioma | G2 | LGG | NaN | NaN | codel | 1 |
| LGG-219 | NaN | NaN | Astrocytoma | G3 | LGG | NaN | NaN | non-codel | 3 |
| LGG-220 | NaN | NaN | Oligodendroglioma | G2 | LGG | NaN | NaN | codel | 1 |
| LGG-223 | NaN | NaN | Oligodendroglioma | G3 | LGG | NaN | NaN | codel | 1 |
| LGG-225 | NaN | NaN | Oligoastrocytoma | G2 | LGG | NaN | NaN | codel | 2 |
| LGG-229 | NaN | NaN | Oligodendroglioma | G2 | LGG | NaN | NaN | codel | 2 |
| LGG-231 | NaN | NaN | Oligoastrocytoma | G3 | LGG | NaN | NaN | codel | 1 |
| LGG-233 | NaN | NaN | Oligodendroglioma | G2 | LGG | NaN | NaN | codel | 3 |
| LGG-234 | NaN | NaN | Oligodendroglioma | G3 | LGG | NaN | NaN | non-codel | 3 |
| LGG-240 | NaN | NaN | Astrocytoma | G3 | LGG | NaN | NaN | non-codel | 3 |
| LGG-241 | NaN | NaN | Oligoastrocytoma | G2 | LGG | NaN | NaN | non-codel | 2 |
| LGG-246 | NaN | NaN | Oligoastrocytoma | G3 | LGG | NaN | NaN | codel | 3 |
| LGG-249 | NaN | NaN | Oligodendroglioma | G2 | LGG | NaN | NaN | codel | 3 |
| LGG-254 | NaN | NaN | Oligodendroglioma | G3 | LGG | NaN | NaN | codel | 2 |
| LGG-260 | NaN | NaN | Oligodendroglioma | G3 | LGG | NaN | NaN | codel | 1 |
| LGG-261 | NaN | NaN | Oligodendroglioma | G2 | LGG | NaN | NaN | codel | 1 |
| LGG-263 | NaN | NaN | Oligoastrocytoma | G3 | LGG | NaN | NaN | non-codel | 3 |
| LGG-269 | NaN | NaN | Oligodendroglioma | G3 | LGG | NaN | NaN | codel | 3 |
| LGG-273 | NaN | NaN | Oligoastrocytoma | G3 | LGG | NaN | NaN | non-codel | 1 |
| LGG-274 | NaN | NaN | Oligoastrocytoma | G2 | LGG | NaN | NaN | codel | 2 |
| LGG-277 | NaN | NaN | Astrocytoma | G2 | LGG | NaN | NaN | non-codel | 3 |
| LGG-278 | NaN | NaN | Oligoastrocytoma | G2 | LGG | NaN | NaN | codel | 2 |
| LGG-280 | NaN | NaN | Oligoastrocytoma | G2 | LGG | NaN | NaN | non-codel | 3 |
| LGG-282 | NaN | NaN | Oligoastrocytoma | G3 | LGG | NaN | NaN | codel | 2 |
| LGG-285 | NaN | NaN | Oligoastrocytoma | G2 | LGG | NaN | NaN | non-codel | 2 |
| LGG-286 | NaN | NaN | Oligodendroglioma | G2 | LGG | NaN | NaN | non-codel | 1 |
| LGG-288 | NaN | NaN | Oligoastrocytoma | G3 | LGG | NaN | NaN | codel | 1 |
| LGG-289 | NaN | NaN | Oligodendroglioma | G2 | LGG | NaN | NaN | codel | 2 |
| LGG-293 | NaN | NaN | Oligoastrocytoma | G2 | LGG | NaN | NaN | non-codel | 3 |
| LGG-295 | NaN | NaN | Oligoastrocytoma | G3 | LGG | NaN | NaN | codel | 3 |
| LGG-296 | NaN | NaN | Oligodendroglioma | G2 | LGG | NaN | NaN | codel | 2 |
| LGG-297 | NaN | NaN | Oligoastrocytoma | G2 | LGG | NaN | NaN | non-codel | 3 |
| LGG-298 | NaN | NaN | Oligoastrocytoma | G3 | LGG | NaN | NaN | codel | 3 |
| LGG-303 | NaN | NaN | Oligodendroglioma | G2 | LGG | NaN | NaN | codel | 1 |
| LGG-304 | NaN | NaN | Oligodendroglioma | G2 | LGG | NaN | NaN | codel | 3 |
| LGG-305 | NaN | NaN | Oligodendroglioma | G2 | LGG | NaN | NaN | codel | 3 |
| LGG-306 | NaN | NaN | Astrocytoma | G3 | LGG | NaN | NaN | non-codel | 3 |
| LGG-307 | NaN | NaN | Oligoastrocytoma | G3 | LGG | NaN | NaN | codel | 2 |
| LGG-308 | NaN | NaN | Oligodendroglioma | G2 | LGG | NaN | NaN | codel | 3 |
| LGG-310 | NaN | NaN | Oligoastrocytoma | G2 | LGG | NaN | NaN | codel | 2 |
| LGG-311 | NaN | NaN | Astrocytoma | G2 | LGG | NaN | NaN | non-codel | 3 |
| LGG-313 | NaN | NaN | Oligoastrocytoma | G2 | LGG | NaN | NaN | non-codel | 2 |
| LGG-314 | NaN | NaN | Oligoastrocytoma | G2 | LGG | NaN | NaN | non-codel | 1 |
| LGG-315 | NaN | NaN | Oligodendroglioma | G2 | LGG | NaN | NaN | codel | 2 |
| LGG-316 | NaN | NaN | Oligodendroglioma | G3 | LGG | NaN | NaN | codel | 1 |
| LGG-320 | NaN | NaN | Oligoastrocytoma | G2 | LGG | NaN | NaN | codel | 1 |
| LGG-321 | NaN | NaN | Oligoastrocytoma | G2 | LGG | NaN | NaN | non-codel | 2 |
| LGG-325 | NaN | NaN | Oligoastrocytoma | G2 | LGG | NaN | NaN | codel | 3 |
| LGG-326 | NaN | NaN | Oligoastrocytoma | G2 | LGG | NaN | NaN | codel | 1 |
| LGG-327 | NaN | NaN | Oligoastrocytoma | G2 | LGG | NaN | NaN | non-codel | 3 |
| LGG-330 | NaN | NaN | Oligoastrocytoma | G3 | LGG | NaN | NaN | codel | 3 |
| LGG-331 | NaN | NaN | Oligodendroglioma | G2 | LGG | NaN | NaN | codel | 2 |
| LGG-333 | NaN | NaN | Oligoastrocytoma | G3 | LGG | NaN | NaN | codel | 2 |
| LGG-334 | NaN | NaN | Oligoastrocytoma | G2 | LGG | NaN | NaN | non-codel | 2 |
| LGG-337 | NaN | NaN | Oligoastrocytoma | G2 | LGG | NaN | NaN | codel | 1 |
| LGG-338 | NaN | NaN | Oligoastrocytoma | G3 | LGG | NaN | NaN | non-codel | 3 |
| LGG-341 | NaN | NaN | Oligoastrocytoma | G3 | LGG | NaN | NaN | codel | 2 |
| LGG-343 | NaN | NaN | Oligoastrocytoma | G2 | LGG | NaN | NaN | non-codel | 2 |
| LGG-344 | NaN | NaN | Oligodendroglioma | G2 | LGG | NaN | NaN | codel | 1 |
| LGG-345 | NaN | NaN | Oligoastrocytoma | G3 | LGG | NaN | NaN | codel | 3 |
| LGG-346 | NaN | NaN | Astrocytoma | G3 | LGG | NaN | NaN | non-codel | 2 |
| LGG-348 | NaN | NaN | Oligoastrocytoma | G2 | LGG | NaN | NaN | codel | 1 |
| LGG-350 | NaN | NaN | Oligoastrocytoma | G2 | LGG | NaN | NaN | codel | 1 |
| LGG-351 | NaN | NaN | Oligoastrocytoma | G2 | LGG | NaN | NaN | non-codel | 1 |
| LGG-352 | NaN | NaN | Oligodendroglioma | G2 | LGG | NaN | NaN | codel | 2 |
| LGG-354 | NaN | NaN | Oligoastrocytoma | G3 | LGG | NaN | NaN | non-codel | 1 |
| LGG-355 | NaN | NaN | Astrocytoma | G3 | LGG | NaN | NaN | codel | 1 |
| LGG-357 | NaN | NaN | Oligoastrocytoma | G2 | LGG | NaN | NaN | codel | 1 |
| LGG-359 | NaN | NaN | Oligodendroglioma | G2 | LGG | NaN | NaN | codel | 3 |
| LGG-360 | NaN | NaN | Oligodendroglioma | G2 | LGG | NaN | NaN | codel | 3 |
| LGG-361 | NaN | NaN | Oligoastrocytoma | G2 | LGG | NaN | NaN | codel | 1 |
| LGG-363 | NaN | NaN | Oligoastrocytoma | G2 | LGG | NaN | NaN | non-codel | 2 |
| LGG-365 | NaN | NaN | Oligoastrocytoma | G3 | LGG | NaN | NaN | codel | 3 |
| LGG-367 | NaN | NaN | Oligoastrocytoma | G2 | LGG | NaN | NaN | codel | 3 |
| LGG-371 | NaN | NaN | Astrocytoma | G3 | LGG | NaN | NaN | non-codel | 1 |
| LGG-373 | NaN | NaN | Oligoastrocytoma | G3 | LGG | NaN | NaN | codel | 2 |
| LGG-374 | NaN | NaN | Oligoastrocytoma | G2 | LGG | NaN | NaN | non-codel | 3 |
| LGG-375 | NaN | NaN | Oligoastrocytoma | G2 | LGG | NaN | NaN | non-codel | 2 |
| LGG-377 | NaN | NaN | Oligodendroglioma | G3 | LGG | NaN | NaN | codel | 1 |
| LGG-380 | NaN | NaN | Oligoastrocytoma | G3 | LGG | NaN | NaN | codel | 1 |
| LGG-383 | NaN | NaN | Oligoastrocytoma | G2 | LGG | NaN | NaN | codel | 3 |
| LGG-385 | NaN | NaN | Oligoastrocytoma | G3 | LGG | NaN | NaN | codel | 2 |
| LGG-387 | NaN | NaN | Oligoastrocytoma | G3 | LGG | NaN | NaN | codel | 2 |
| LGG-388 | NaN | NaN | Oligoastrocytoma | G2 | LGG | NaN | NaN | codel | 1 |
| LGG-391 | NaN | NaN | Oligoastrocytoma | G2 | LGG | NaN | NaN | non-codel | 1 |
| LGG-394 | NaN | NaN | Oligodendroglioma | G3 | LGG | NaN | NaN | codel | 3 |
| LGG-395 | NaN | NaN | Oligoastrocytoma | G3 | LGG | NaN | NaN | codel | 1 |
| LGG-396 | NaN | NaN | Oligoastrocytoma | G2 | LGG | NaN | NaN | codel | 1 |
| LGG-492 | NaN | NaN | Oligoastrocytoma | G2 | LGG | NaN | NaN | codel | 2 |
| LGG-500 | NaN | NaN | Oligoastrocytoma | G2 | LGG | NaN | NaN | non-codel | 1 |
| LGG-506 | NaN | NaN | Oligoastrocytoma | G2 | LGG | NaN | NaN | non-codel | 1 |
| LGG-515 | NaN | NaN | Oligoastrocytoma | G2 | LGG | NaN | NaN | codel | 3 |
| LGG-516 | NaN | NaN | Oligoastrocytoma | G3 | LGG | NaN | NaN | non-codel | 2 |
| LGG-518 | NaN | NaN | Astrocytoma | G3 | LGG | NaN | NaN | non-codel | 3 |
| LGG-519 | NaN | NaN | Oligoastrocytoma | G2 | LGG | NaN | NaN | non-codel | 1 |
| LGG-520 | NaN | NaN | Oligodendroglioma | G2 | LGG | NaN | NaN | codel | 2 |
| LGG-525 | NaN | NaN | Oligodendroglioma | G2 | LGG | NaN | NaN | codel | 2 |
| LGG-527 | NaN | NaN | Oligoastrocytoma | G2 | LGG | NaN | NaN | codel | 3 |
| LGG-532 | NaN | NaN | Oligoastrocytoma | G3 | LGG | NaN | NaN | non-codel | 3 |
| LGG-533 | NaN | NaN | Oligoastrocytoma | G2 | LGG | NaN | NaN | non-codel | 1 |
| LGG-537 | NaN | NaN | Oligoastrocytoma | G2 | LGG | NaN | NaN | non-codel | 2 |
| LGG-545 | NaN | NaN | Oligoastrocytoma | G2 | LGG | NaN | NaN | non-codel | 1 |
| LGG-547 | NaN | NaN | Oligodendroglioma | G2 | LGG | NaN | NaN | codel | 2 |
| LGG-550 | NaN | NaN | Oligoastrocytoma | G3 | LGG | NaN | NaN | codel | 3 |
| LGG-552 | NaN | NaN | Astrocytoma | G3 | LGG | NaN | NaN | non-codel | 2 |
| LGG-558 | NaN | NaN | Oligoastrocytoma | G2 | LGG | NaN | NaN | non-codel | 1 |
| LGG-561 | NaN | NaN | Oligoastrocytoma | G3 | LGG | NaN | NaN | codel | 2 |
| LGG-563 | NaN | NaN | Oligodendroglioma | G2 | LGG | NaN | NaN | codel | 2 |
| LGG-565 | NaN | NaN | Oligoastrocytoma | G2 | LGG | NaN | NaN | codel | 3 |
| LGG-566 | NaN | NaN | Oligoastrocytoma | G2 | LGG | NaN | NaN | codel | 3 |
| LGG-570 | NaN | NaN | Oligoastrocytoma | G2 | LGG | NaN | NaN | codel | 2 |
| LGG-572 | NaN | NaN | Astrocytoma | G3 | LGG | NaN | NaN | codel | 3 |
| LGG-573 | NaN | NaN | Oligoastrocytoma | G2 | LGG | NaN | NaN | codel | 1 |
| LGG-574 | NaN | NaN | Oligodendroglioma | G2 | LGG | NaN | NaN | non-codel | 1 |
| LGG-576 | NaN | NaN | Oligoastrocytoma | G2 | LGG | NaN | NaN | codel | 1 |
| LGG-579 | NaN | NaN | Oligoastrocytoma | G2 | LGG | NaN | NaN | codel | 1 |
| LGG-581 | NaN | NaN | Oligoastrocytoma | G2 | LGG | NaN | NaN | codel | 2 |
| LGG-582 | NaN | NaN | Oligodendroglioma | G2 | LGG | NaN | NaN | codel | 2 |
| LGG-585 | NaN | NaN | Astrocytoma | G2 | LGG | NaN | NaN | non-codel | 3 |
| LGG-587 | NaN | NaN | Oligodendroglioma | G3 | LGG | NaN | NaN | codel | 3 |
| LGG-589 | NaN | NaN | Oligoastrocytoma | G2 | LGG | NaN | NaN | non-codel | 2 |
| LGG-590 | NaN | NaN | Oligoastrocytoma | G2 | LGG | NaN | NaN | codel | 1 |
| LGG-591 | NaN | NaN | Astrocytoma | G3 | LGG | NaN | NaN | non-codel | 1 |
| LGG-593 | NaN | NaN | Oligodendroglioma | G3 | LGG | NaN | NaN | codel | 1 |
| LGG-594 | NaN | NaN | Oligoastrocytoma | G3 | LGG | NaN | NaN | non-codel | 2 |
| LGG-597 | NaN | NaN | Oligoastrocytoma | G2 | LGG | NaN | NaN | codel | 1 |
| LGG-600 | NaN | NaN | Oligoastrocytoma | G3 | LGG | NaN | NaN | codel | 3 |
| LGG-601 | NaN | NaN | Astrocytoma | G3 | LGG | NaN | NaN | non-codel | 3 |
| LGG-604 | NaN | NaN | Oligoastrocytoma | G2 | LGG | NaN | NaN | codel | 2 |
| LGG-607 | NaN | NaN | Oligodendroglioma | G2 | LGG | NaN | NaN | codel | 3 |
| LGG-609 | NaN | NaN | Oligoastrocytoma | G2 | LGG | NaN | NaN | non-codel | 2 |
| LGG-610 | NaN | NaN | Oligoastrocytoma | G2 | LGG | NaN | NaN | non-codel | 3 |
| LGG-612 | NaN | NaN | Oligodendroglioma | G2 | LGG | NaN | NaN | codel | 2 |
| LGG-613 | NaN | NaN | Oligoastrocytoma | G2 | LGG | NaN | NaN | non-codel | 1 |
| LGG-614 | NaN | NaN | Oligodendroglioma | G2 | LGG | NaN | NaN | codel | 3 |
| LGG-616 | NaN | NaN | Oligoastrocytoma | G2 | LGG | NaN | NaN | codel | 3 |
| LGG-620 | NaN | NaN | Oligodendroglioma | G2 | LGG | NaN | NaN | codel | 3 |
| LGG-622 | NaN | NaN | Oligoastrocytoma | G2 | LGG | NaN | NaN | non-codel | 2 |
| LGG-624 | NaN | NaN | Oligoastrocytoma | G3 | LGG | NaN | NaN | non-codel | 2 |
| LGG-625 | NaN | NaN | Oligoastrocytoma | G2 | LGG | NaN | NaN | non-codel | 2 |
| LGG-626 | NaN | NaN | Astrocytoma | G3 | LGG | NaN | NaN | codel | 2 |
| LGG-630 | NaN | NaN | Oligodendroglioma | G2 | LGG | NaN | NaN | codel | 3 |
| LGG-631 | NaN | NaN | Oligoastrocytoma | G2 | LGG | NaN | NaN | non-codel | 1 |
| LGG-632 | NaN | NaN | Oligoastrocytoma | G2 | LGG | NaN | NaN | codel | 2 |
| LGG-634 | NaN | NaN | Oligodendroglioma | G3 | LGG | NaN | NaN | codel | 2 |
| LGG-637 | NaN | NaN | Oligodendroglioma | G2 | LGG | NaN | NaN | codel | 1 |
| LGG-639 | NaN | NaN | Oligoastrocytoma | G2 | LGG | NaN | NaN | codel | 1 |
| LGG-642 | NaN | NaN | Oligodendroglioma | G3 | LGG | NaN | NaN | codel | 2 |
| LGG-647 | NaN | NaN | Oligoastrocytoma | G2 | LGG | NaN | NaN | non-codel | 3 |
| LGG-648 | NaN | NaN | Astrocytoma | G2 | LGG | NaN | NaN | codel | 3 |
| LGG-651 | NaN | NaN | Oligodendroglioma | G2 | LGG | NaN | NaN | codel | 3 |
| LGG-658 | NaN | NaN | Oligodendroglioma | G3 | LGG | NaN | NaN | codel | 3 |
| LGG-659 | NaN | NaN | Oligoastrocytoma | G2 | LGG | NaN | NaN | codel | 2 |
| LGG-660 | NaN | NaN | Oligoastrocytoma | G2 | LGG | NaN | NaN | codel | 1 |
| LGG-766 | NaN | NaN | Oligoastrocytoma | G2 | LGG | NaN | NaN | non-codel | 1 |
